# Single nucleus transcriptome analysis of *Arabidopsis thaliana* roots infected with *Phytophthora. capsici*

**DOI:** 10.64898/2026.04.28.720150

**Authors:** Razan M Alajoleen, Tran N. Chau, Joel Shuman, Bastiaan O.R. Bargmann, Song Li

**Author notes:** Corresponding Author: Song Li.

## Abstract

Understanding how plant roots coordinate immune responses at the cellular level is key to unraveling host–pathogen interactions. Using single-nucleus RNA sequencing (snRNA-seq) of *Arabidopsis thaliana* roots 24 hours after *Phytophthora. capsici*(*P. capsici*)inoculation, we captured the transcriptional landscape of early infection at single-cell resolution. Four libraries (two infected and two mock-treated) were generated with approximately 26,000 high-quality nuclei with consistent sequencing depth and viability. A reference-based pipeline distinguished host and pathogen transcripts, enabling species-resolved mapping and host-focused single-nucleus transcriptomic analysis. Integration and clustering identified 12 transcriptionally distinct root cell types, encompassing major tissues such as the meristem, cortex, endodermis, and vasculature. Cluster-specific marker analysis confirmed cell-type identities, while differential expression and Gene Ontology enrichment revealed a global transcriptional shift from metabolic and translational processes in mock samples to defense-, stress-, and pathogen-response pathways upon infection. Hormone-related enrichment indicated broad salicylic acid activation across root tissues, spatially confined ethylene signaling in vascular-associated clusters, and localized jasmonic acid responses in cortex and phloem. Together, these results provide a high-resolution view of *Arabidopsis* root immunity, highlighting a coordinated yet tissue-specific defense architecture in which salicylic acid underpins systemic protection, ethylene modulates vascular defense, and jasmonic acid contributes targeted reinforcement during early *P. capsici* infection.

## Introduction

Oomycetes are a diverse group of fungus-like eukaryotes that cause severe diseases in plants. Unlike true fungi, they belong to the Stramenopiles, a lineage that includes brown algae and diatoms, and exhibit distinct evolutionary and cellular features[1]. Within this group, the genus *Phytophthora* comprises some of the most destructive plant pathogens worldwide [2]. *P. capsici* is particularly damaging due to its broad host range and aggressive infection strategy, causing yield losses exceeding 90% under favorable conditions [3, 4].

*P. capsici* is a hemibiotrophic pathogen that infects multiple plant organs and transitions from an initial biotrophic phase to a necrotrophic phase involving extensive tissue damage [5, 6]. This lifestyle transition is accompanied by extensive host transcriptional and cellular reprogramming[7] and is driven in part by pathogen-secreted effector proteins that suppress host immunity and promote colonization [8]. Despite its agricultural importance, the molecular basis of *P. capsici*–host interactions remains incompletely understood, particularly at early infection stages, due in part to limited genetic resources in many crop hosts [9].

*Arabidopsis thaliana* provides a powerful model for dissecting conserved mechanisms of plant–oomycete interactions, owing to its well-characterized immune system and extensive genetic and genomic resources [10]. Although *P. capsici* is not a natural pathogen of *Arabidopsis*, it can infect *Arabidopsis* roots under controlled conditions, enabling detailed investigation of cell type-specific defense responses [4]. Previous studies have identified roles for salicylic acid signaling, reactive oxygen species, and defense gene activation in *Arabidopsis* responses to *P. capsici*[11]; however, the cellular heterogeneity of root tissues has limited resolution of these responses.

Recent advances in single-cell and single-nucleus RNA sequencing have enabled high-resolution analysis of plant immune responses at the level of individual cell types [12]. Applying these approaches to the *Arabidopsis*–*P. capsici* pathosystem provides a unique opportunity to resolve spatially and temporally distinct transcriptional responses during early infection, particularly at 24 hours post-inoculation, when host reprogramming is most active.

We used single-nucleus RNA sequencing (snRNA-seq) to profile *Arabidopsis* root responses to *P. capsici* infection at 24 hours post-inoculation. By integrating infected and non-infected datasets, we identified transcriptionally distinct root cell populations and defined cell type-specific defense programs involving hormone signaling, oxidative stress responses, and chromatin regulation. Together, these results provide a high-resolution view of early root immune responses and reflect **Dr. McDowell’s** enduring commitment to elucidating the cellular mechanisms underlying plant immunity.

## Results

### snRNA-seq reveals cell type-specific transcriptional profiles of *Arabidopsis* roots infected with *P. capsici*

To investigate cell-type-specific transcriptional reprogramming during early infection, we infected 7-day-old *Arabidopsis* (Col-0) roots with *P. capsici*, while mock roots were inoculated with buffered nodulation medium (BNM) as controls. 24 hours post-inoculation (hpi), nuclei were isolated from infected and noninfected roots and processed for snRNA-seq using the 10x Genomics Chromium platform (*Fig. 1a*). A total of 26,114 high-quality single-nucleus transcriptomes were recovered from four libraries (two infected and two mock), with a median of 549 genes and 974 unique molecular identifiers (UMIs) detected per nucleus *(Supplementary Fig. 1.a*).

**Figure 1.**
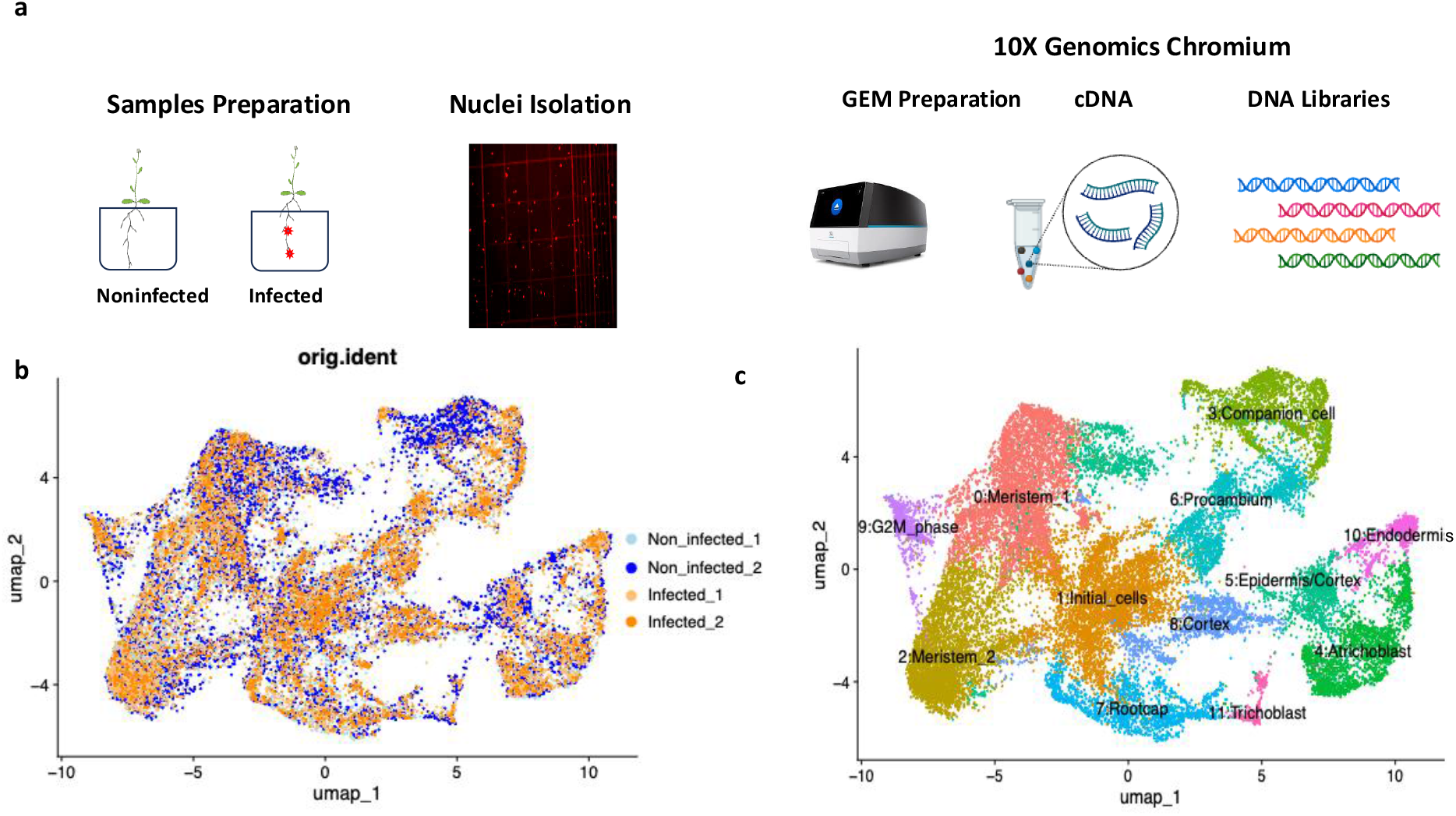
Single-nucleus RNA-seq (snRNA-seq) transcriptomic profiling of *Arabidopsis thaliana* roots infected with *P. capsici*. (a) Overview of the experimental and analytical workflow. Roots from *Arabidopsis thaliana* Col-0 plants were collected at 24 h post-inoculation (hpi) with *P. capsici* (Infected) and mock-treated controls (Non_Infected). Nuclei were isolated from frozen root tissue, stained with DAPI, and sorted prior to encapsulation using the 10x Genomics Chromium platform. Single nuclei were partitioned into Gel Beads-in-Emulsion (GEMs) for reverse transcription and barcoding, followed by cDNA amplification and construction of indexed DNA libraries for sequencing. (b) UMAP visualization of integrated nuclei datasets from four biological replicates (Non_Infected_1, Non_Infected_2, Infected_1, Infected_2). Each replicate is color-coded, showing effective dataset alignment and minimal batch variation after integration. (c) Cell-type clustering of the integrated dataset, revealing transcriptionally distinct populations corresponding to known *Arabidopsis* root tissues, including meristematic zone, cortex, endodermis, procambium, and root cap. Workflow adapted from 10x Genomics Single Cell Gene Expression v3 chemistry (10x Genomics, 2021).

After quality control filtering (min.features = 200, min.cells = 3) and data integration using Seurat v5.3.0 [13] twelve transcriptionally distinct clusters were identified (*Fig. 1b–c*). Marker genes for each cluster were determined using the *FindMarkers* function (min.pct = 0.01, logfc.threshold = 0.25) and used for cell-type annotation with the Orthologous Marker Gene (OMG) framework [14] (*Supplementary Fig. 2)*. This analysis resolved major *Arabidopsis* root tissues, including meristematic, cortical, endodermal, procambium, and root cap cell populations, providing a detailed transcriptomic atlas for infection-associated responses at the single-nucleus level.

To characterize transcriptional heterogeneity among root nuclei, we identified marker genes for each of the thirteen clusters using Seurat’s *FindAllMarkers* function (min.pct = 0.01, logfc.threshold = 0.25). Distinct cell type-specific gene signatures were obtained for major root tissues, including the meristematic zone, cortex, endodermis, procambium, and phloem (*Supplementary Fig. 3a*). The top markers included *CASP1* (AT2G36100), *EXPA7* (AT1G12560), *APL* (AT1G79430), and *ANAC033* (AT1G79580), which are well-established indicators of endodermal, root hair, phloem, and procambium cell types, respectively [15].

UMAP feature plots confirmed that these canonical markers were spatially restricted to their corresponding cell populations *(Supplementary Fig. 3b)*. The expression pattern of *CASP1* was confined to endodermal clusters, while *APL* and *ANAC033* were enriched in phloem and procambium-associated cells, respectively, demonstrating accurate cell-type delineation within the dataset.

### snRNA-seq quality assessment and transcriptomic profiling during infection

To ensure the generation of high-quality single-nucleus transcriptomes, we evaluated nuclei yield, viability, and sequencing performance across four *Arabidopsis* root libraries (two infected and two mock). Nuclei isolation produced nuclei densities ranging from 327,500 to 460,000 cells/ml with viabilities between 83.06% and 94.63% (*Fig. 2a*), confirming successful recovery of intact nuclei suitable for snRNA-seq. Sequencing and alignment metrics were consistent with high-quality data across replicates (*Supplementary Fig.1.b*). Each library yielded approximately 8,000 nuclei, with mean reads per cell between 61,091 and 124,296, median genes per nucleus between 158 and 964, and read mapping efficiency ranging from 43.0% to 95.7%. In total, 2.86 × 10^9^ reads were generated using the 10x Genomics Cell Ranger v8.0.1 pipeline.

**Figure 2.**
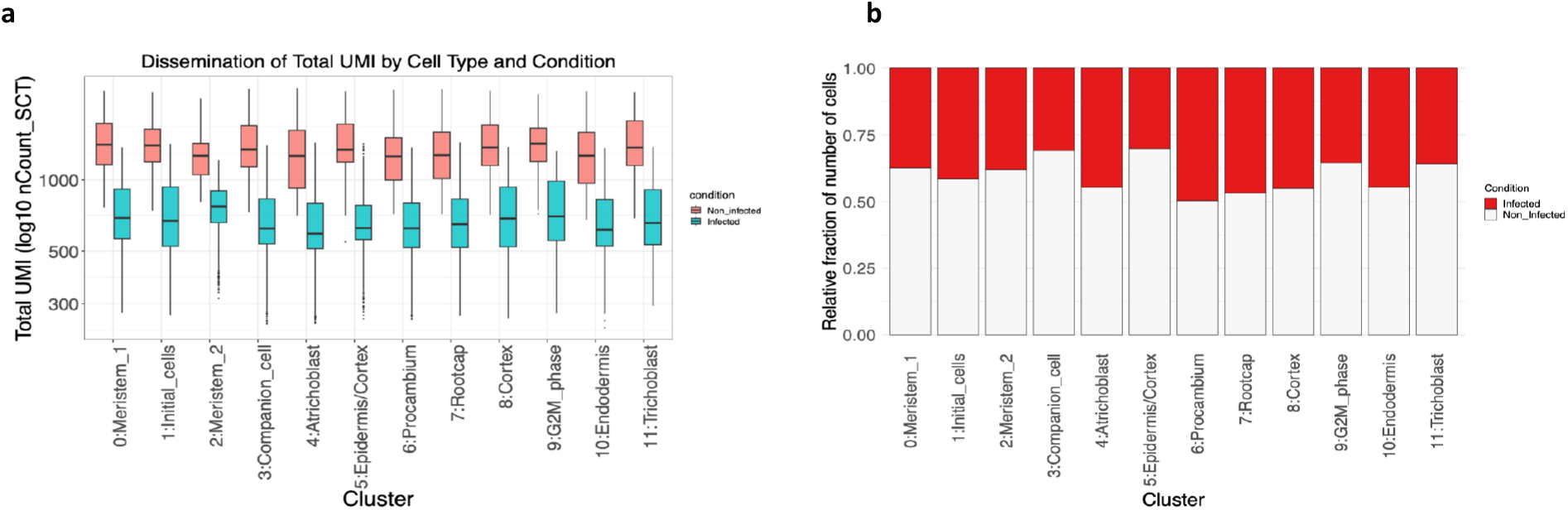
Transcriptomic profiling of *Arabidopsis thaliana* root cells during *P. capsici* infection. (a) Distribution of total unique molecular identifiers (UMIs) across annotated cell types in infected and noninfected roots. Infected roots showed slightly higher transcript abundance in specific cell types, suggesting condition-specific transcriptional activation. (b) Relative proportion of nuclei per annotated cluster across conditions. The fraction of nuclei from infected roots increased notably in clusters associated with defense- and hormone-related transcriptional programs, reflecting infection-driven changes in cell-type composition. Sequencing reads were processed using the Cell Ranger v8.0.1 pipeline (10x Genomics, 2023), and gene-barcode matrices were imported into Seurat v5 (Stuart et al., 2019) for downstream analysis.

Comparison of total unique molecular identifiers (UMIs) by cell type revealed cell type-specific transcriptional variation between infected and control samples (*Fig. 2a*). Differences in cell-type proportions between infected and non-infected samples likely reflect cell type–dependent effects of infection on nuclei recovery and representation and should therefore be interpreted cautiously. For example, procambium, root cap cluster 2, and cortex cells all showed an increased proportion following infection compared with other cell clusters (*Fig. 2b*).

Collectively, these results confirm the robustness and reproducibility of the snRNA-seq dataset and provide a foundation for exploring the transcriptional dynamics underlying *Arabidopsis–Phytophthora* interactions at single-nucleus resolution.

### Gene Ontology enrichment reveals infection-specific biological processes

To elucidate the functional pathways underlying cell type-specific transcriptional reprogramming during infection, we performed Gene Ontology (GO) enrichment analysis on cluster-specific marker genes derived from infected and noninfected roots. Differentially expressed genes were obtained using Seurat’s FindMarkers function (min.pct = 0.01, logfc.threshold = 0.25) and GO Biological Process enrichment was conducted using the clusterProfiler package (v4.0; Wu et al., 2021) with p.adjust < 0.05 (Benjamini–Hochberg correction).

The GO enrichment analysis revealed distinct biological processes between infected and non-infected nuclei across the major root cell clusters (*Fig. 3*). Infected nuclei showed significant enrichment of stress- and metabolism-associated pathways, including *cellular response to oxidative stress, response to hypoxia*, and *glycolytic process*, suggesting activation of redox regulation and energy reprogramming during infection. In contrast, non-infected nuclei were enriched for fundamental biosynthetic and metabolic processes such as *translation, RNA splicing*, and *ATP synthesis coupled electron transport*, reflecting active protein synthesis and mitochondrial function under normal physiological conditions. The distribution of GO terms across clusters indicates that infection triggers broad cellular adjustments, with meristematic and cortical clusters exhibiting the strongest transcriptional responses.

**Figure 3.**
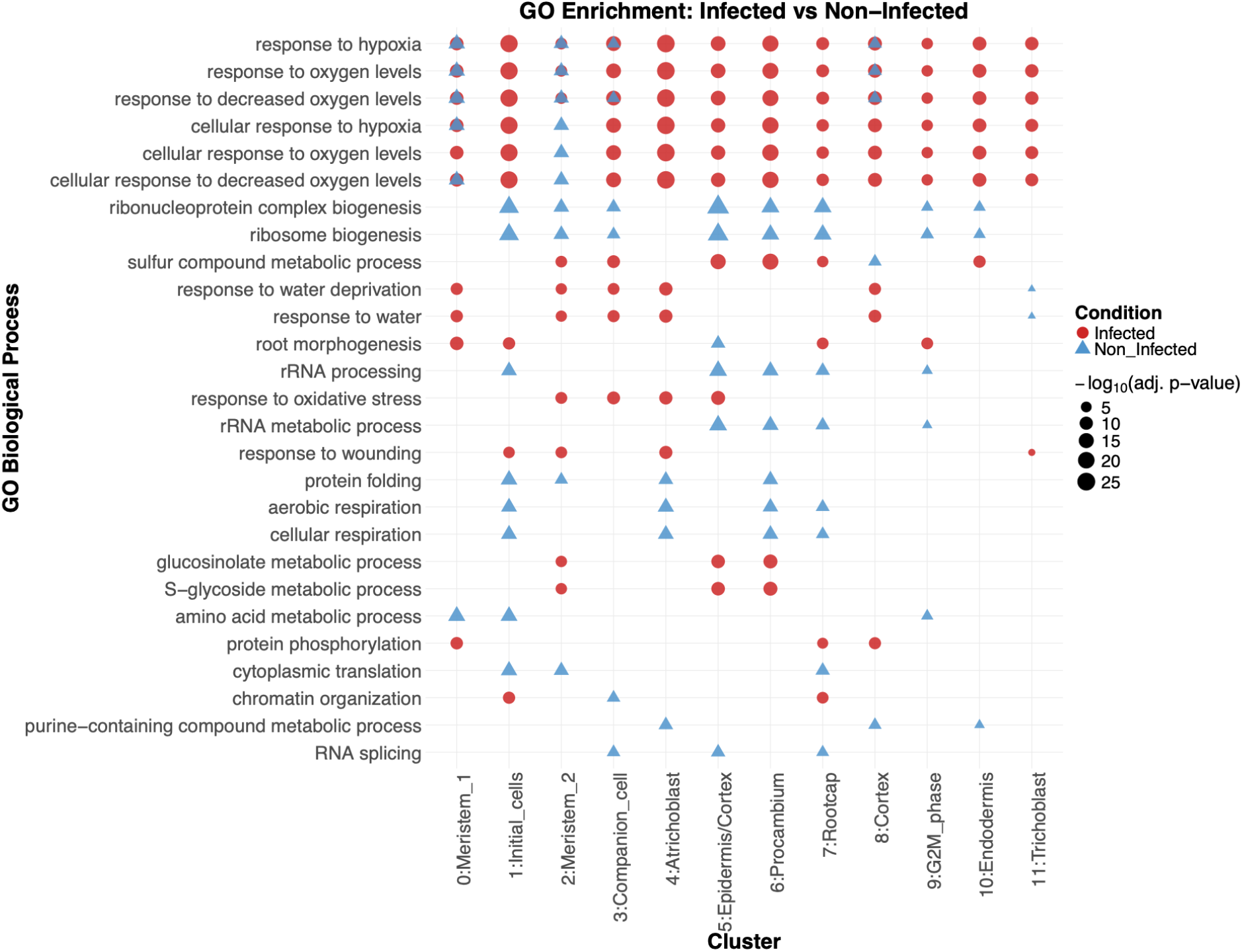
Dot plot of Gene Ontology (GO) enrichment in *Arabidopsis thaliana* root nuclei during *P. capsici* infection. Dot plot summarizing significantly enriched Gene Ontology (GO) Biological Process terms across twelve cell clusters of *Arabidopsis thaliana* root nuclei under infected (red) and noninfected (blue) conditions. The size of each dot corresponds to the statistical significance of enrichment (−log_10_ adjusted p-value), and the shape represents the experimental condition. GO enrichment was calculated using the clusterProfiler R package (Wu et al., 2021) with Benjamini-Hochberg correction (p < 0.05).

### Hormone signaling and defense activation during *P. capsici* infection

To resolve hormone signaling underlying the *Arabidopsis* root response to *P. capsici*, we performed cluster-resolved GO enrichment on hormone-related Biological Process terms using DEGs between infected and non-infected roots. In infected roots, salicylic acid terms-*salicylic acid mediated signaling pathway, cellular response to salicylic acid stimulus*, and *regulation of salicylic acid mediated signaling pathway*-were significantly enriched across meristematic (cluster 0), cortex (8), endodermis (10), and trichoblasts (11) relative to controls (FDR < 0.05), indicating broad SA pathway activation early after infection. In contrast, ethylene terms-*ethylene-activated signaling pathway* and *cellular response to ethylene stimulus*-were predominantly enriched in procambium-associated clusters (6 and 9), consistent with localized ET signaling. Jasmonic acid (JA) terms-jasmonic acid mediated signaling pathway and cellular response to jasmonic acid stimulus-showed a moderate but significant enrichment in cortex (8) and phloem (3), suggesting a supportive, tissue-specific contribution of JA. Additional hormone categories (abscisic acid, auxin, brassinosteroid, cytokinin, gibberellin) were detected but were weaker and/or cluster-restricted. Collectively, these enrichments support a model in which SA and ET dominate the early immune response, with JA reinforcing defense in select tissues (*Fig. 4a*).

**Figure 4.**
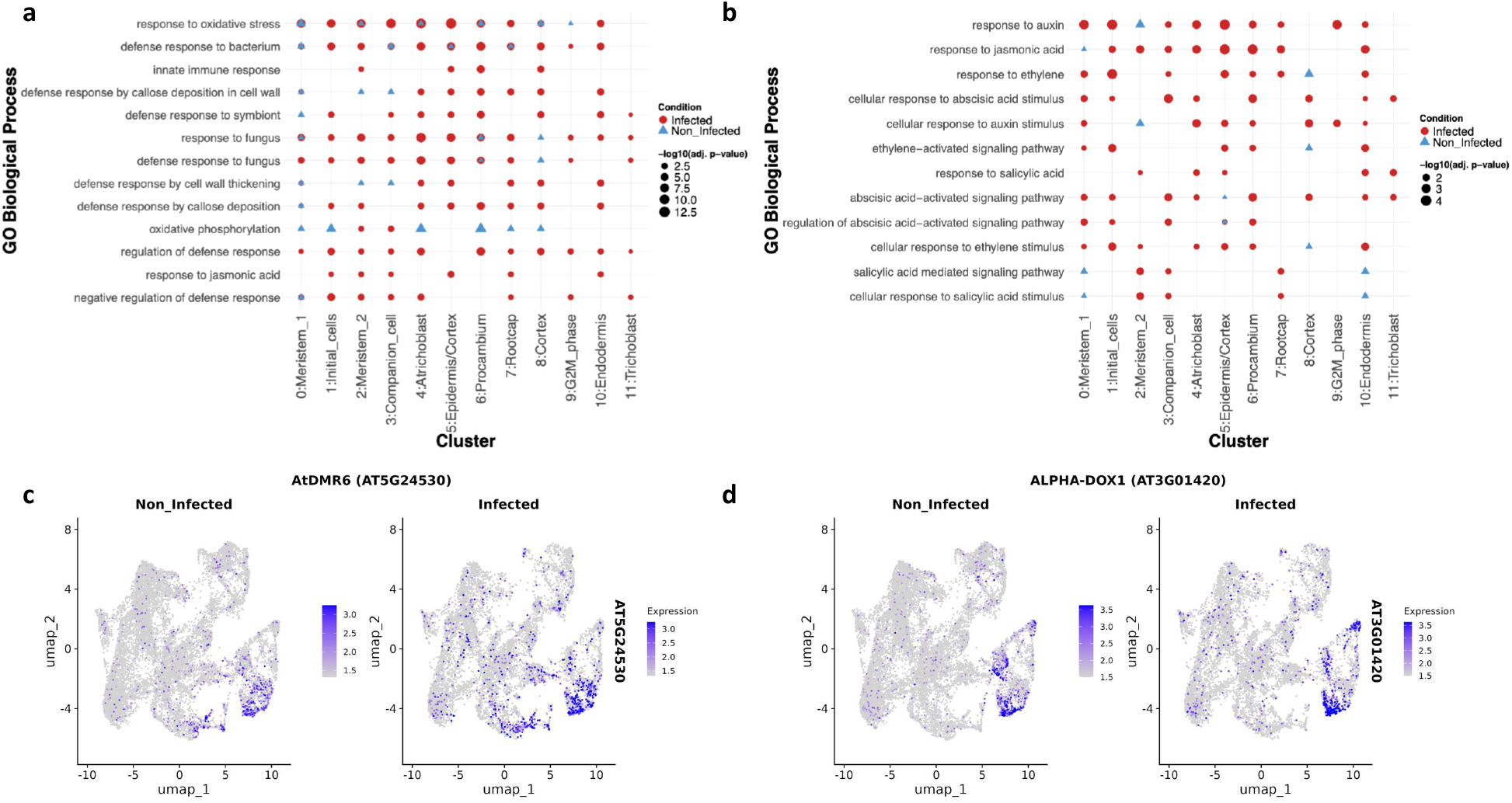
Cluster-resolved GO enrichment and marker gene expression in *Arabidopsis* roots responding to *P. capsici*. (a) Hormone response GO terms. Bubble plot of GO Biological Process terms related to plant hormones across cell clusters (x-axis; cluster ID and inferred label). Each symbol represents the condition (red circles = Infected, blue triangles = Non_Infected). Bubble size scales with –log10(adjusted p-value) from over-representation testing (Benjamini-Hochberg correction). Terms include salicylic acid (SA), ethylene (ET), jasmonic acid (JA), abscisic acid (ABA), auxin, brassinosteroid, and cytokinin pathways. (b) Defense and oomycete/innate immunity GO terms. Same layout and statistics as in (a), showing enrichment for defense, immune system processes, responses to oomycete/fungus/bacterium/virus, SA signaling/ metabolism, oxidative stress, and related categories. (c) Gene-level expression maps. UMAP feature plots for representative defense-associated genes shown separately for Non_Infected and Infected cells: AtDMR6 (AT5G24530) (left pair) and ALPHA-DOX1 (AT3G01420) (right pair). Color bar indicates normalized expression (lighter to darker blue = higher expression). Statistics: GO enrichment was performed per cluster by comparing Infected vs Non_Infected differentially expressed genes; p-values were adjusted by Benjamini–Hochberg. Abbreviations: SA, salicylic acid; ET, ethylene; JA, jasmonic acid; ABA, abscisic acid.

To contextualize these transcriptional changes, we performed GO enrichment analysis of differentially expressed genes between infected and noninfected roots. Infected clusters exhibited strong enrichment for defense-related processes such as innate immune response, response to jasmic acid, immune system process, and regulation of defense response (*Fig. 4b*). In contrast, noninfected clusters were dominated by GO terms associated with basal metabolic and translational processes, consistent with a non-stressed physiological state.

UMAP visualization of representative defense genes supported these findings. *AtDMR6* (AT5G24530), encoding a salicylic acid 3-hydroxylase, was highly expressed in infected cortex and endodermal nuclei as compared to control, consistent with its role in fine-tuning SA accumulation and immunity [16]. Similarly, *ALPHA-DOX1* (AT3G01420), a lipid-peroxidizing enzyme linked to oxylipin-mediated defense[17] showed increased expression in infected roots (*Fig. 4c*). Together, these results indicate that *P. capsici* infection triggers a coordinated transcriptional reprogramming across multiple hormone pathways.

## Discussion

Early infection represents a critical phase during which *P. capsici* establishes compatibility with *Arabidopsis* roots and initiates host signaling programs that shape disease progression. We selected 24 hours post-infection (hpi) to capture the biotrophic phase, when haustoria are formed and active host–pathogen interactions occur. Cytological studies show that zoospore attachment, penetration, and haustorium formation occur by 24 hpi, whereas sporangia emerge by 48 hpi, marking the transition to necrotrophy [18]. Consistent with this timeline, trypan blue staining in our laboratory revealed haustoriated cortical cells at 24 hpi, supporting this time point as optimal for profiling early transcriptional reprogramming.

Our snRNA-seq analysis resolved 12 *Arabidopsis* root cell clusters and revealed spatially distinct immune and hormone-associated transcriptional programs. GO enrichment analysis showed salicylic acid (SA) signaling as a dominant and broadly enriched pathway across multiple tissues, including meristematic, cortical, and endodermal cells, suggesting a root-wide immune backbone. In contrast, ethylene (ET) signaling was strongly enriched in procambium-associated clusters, consistent with vascular-localized defense responses, while jasmonic acid (JA) and abscisic acid (ABA) pathways displayed tissue-specific enrichment, particularly in cortex and phloem. Limited enrichment of auxin and brassinosteroid pathways further supports cell type-specific hormonal regulation rather than a global transcriptional shift.

SA-regulated genes such as *AtDMR6* were highly expressed in infected cortex and endodermis, indicating localized modulation of SA signaling similar to resistance mechanisms described in tomato (*SlDMR6-1*). Additionally, induction of oxylipin-related genes, including *ALPHA-DOX1*, suggests integration of lipid-derived signaling during early infection [19].

While our analysis revealed pronounced infection-associated transcriptional responses across multiple root cell types, interpretation of apparent changes in cell-type abundance must be approached with caution. Differences in nuclei isolation efficiency between infected and mock samples may influence relative cell-type representation, potentially confounding conclusions regarding infection-induced shifts in cellular composition[20]. Accordingly, rather than asserting definitive remodeling of root cellular architecture, we emphasize infection-associated transcriptional reprogramming within individual cell types. This distinction is important, as technical variability inherent to nuclei recovery can impact proportional analyses.

Together, our findings support a model in which *P. capsici* establishes biotrophic infection by 24 hpi while *Arabidopsi*s roots deploy coordinated, tissue-specific immune responses dominated by SA signaling, refined by ET in vascular tissues, and reinforced by JA and ABA in select cell types. This study highlights 24 hpi as a critical window for dissecting early host–oomycete interactions and demonstrates the power of snRNA-seq for resolving cell-type-specific immune dynamics in plant roots.

## Materials and Methods

### Plant Growth, Inoculation, and Tissue Collection

*Arabidopsis* thaliana Col-0 CORTEX reporter seeds (pCORTEX::GFP), used to visualize root cortex cells, were surface-sterilized using 60 mg dichloroisocyanuric acid sodium salt hydrate per 1 mL deionized water mixed with 1 mL 100% ethanol, shaken at 100 rpm for 20 min, followed by three washes with 100% ethanol. Sterilized seeds were grown in sterile phytatrays (Sigma-Aldrich, Cat. # P5929) containing approximately 250 mL of 0.5× Murashige and Skoog (MS) medium supplemented with 10% (w/v) sucrose.

Approximately 500 seeds were evenly distributed per tray onto a 200 µm food-grade woven nylon mesh (Amazon, Cat. # B0HYHHX5V) placed on pipette tip racks cut to fit the phytatrays, with four 0.5 mL microcentrifuge tubes positioned at each corner for buoyancy. Plants were grown under a 12 h light/12 h dark photoperiod at 25 °C in a Percival incubator (Model CU36L5).

At 9 days post-germination, roots were inoculated with 10 mL of *P. capsici*Ra zoospore suspension (50,000 zoospores/mL). Zoospores were generated using a lab-specific unpublished protocol. Mock-inoculated seedlings served as non-infected controls. Root tissues from infected and mock-treated plants were harvested at 24 h post-inoculation (hpi).

Two experimental conditions were analyzed: infected (infected_1 and infected_2) and non-infected (non_infected_1 and non_infected_2), with two biological replicates per condition. Roots were excised using sterile scalpels and collected into 60 mm Petri dishes containing dry Kimwipes to remove excess liquid. Approximately 25–30 mg of tissue per sample was transferred to pre-chilled 35 mm Petri dishes for nuclei isolation.

### Nuclei Isolation from Root Tissue

Nuclei were isolated using a modified protocol adapted from the Birnbaum lab [21] to preserve transcriptomic integrity. All procedures were performed on ice or at 4 °C unless otherwise stated. Lysis, wash, and final buffers were freshly supplemented with DTT, PMSF, RNase inhibitor, protease inhibitor cocktail, BSA, and DEPC-treated water. Root tissue was finely chopped in lysis buffer on ice and further homogenized using a Dounce homogenizer (10 strokes each with pestles A and B, with an intermediate incubation on ice).

The homogenate was filtered through a 20 µm mesh and centrifuged at 1,000 × g for 10 min at 4 °C. The pellet was washed, recentrifuged, and resuspended in final buffer, followed by filtration through a 10 µm mesh to obtain a clean nuclei suspension.

Nuclei concentration and integrity were assessed by staining with DAPI or propidium iodide and counting on a hemocytometer under fluorescence microscopy. Only intact, fluorescent-positive nuclei were counted. Nuclei concentrations were used to determine optimal loading for the 10x Genomics Chromium platform for single-nucleus RNA sequencing.

### Generation and Analysis of Single-Nucleus Transcriptomes

Single-nucleus RNA-seq (snRNA-seq) libraries were prepared using the Chromium Next GEM Single Cell 3ʹ Reagent Kit v3.1 (10x Genomics, Pleasanton, CA, USA) according to the manufacturer’s instructions. Nuclei suspensions were adjusted to the desired concentration and loaded onto the Chromium Controller to generate Gel Beads-in-Emulsion (GEMs). Within each GEM, nuclei were lysed and polyadenylated transcripts were barcoded with unique cell barcodes and unique molecular identifiers (UMIs) during reverse transcription.

Following cDNA recovery and amplification, libraries were enzymatically fragmented, end-repaired, A-tailed, and ligated to sequencing adapters. Sample indices were added by PCR, and libraries were purified using SPRIselect beads (Beckman Coulter, USA). Library concentration was quantified using a Qubit fluorometric assay (Thermo Fisher Scientific), and fragment size distribution was assessed with a Bioanalyzer High Sensitivity DNA Kit (Agilent Technologies). Libraries were pooled equimolarly and sequenced on an Illumina NovaSeq X Plus PE150 platform to a depth of approximately 50,000–100,000 reads per nucleus, as recommended by 10x Genomics.

### Species-Resolved Mapping Pipeline for Plant-Pathogen Data

To distinguish *Arabidopsis thaliana* and *P. capsici* transcripts in mixed snRNA-seq libraries, a multi-species reference mapping pipeline was implemented (*Supplementary Fig. S4*). Raw BCL files were processed using Cell Ranger v8.0.1 with a combined *A. thaliana* (TAIR10) and *P. capsici* reference genome. The Cell Ranger “multi” pipeline generated a gem_classification.csv file, which was used to classify barcodes as plant- or pathogen-derived.

Species-specific barcodes were extracted and used with the subset-bam tool to generate species-resolved BAM files, which were converted to FASTQ format using bamtofastq. FASTQ files were then aligned independently to their respective reference genomes using Cell Ranger count, generating separate expression matrices for *Arabidopsis* and *P. capsici*. This approach minimized cross-mapping and enabled downstream differential expression, gene ontology (GO), and pathway analyses. All commands and parameters are detailed in Supplementary Fig. S2.

### Data Integration and Clustering

Downstream analyses were performed in R version 4.4.2 using Seurat v5.3.0 [13]. Count matrices were imported using Read10X(), filtered (min.features = 200; min.cells = 3), and normalized using LogNormalize(). The top 2,000 highly variable features were identified with FindVariableFeatures(). Integration anchors were computed using FindIntegrationAnchors(), followed by dataset integration using IntegrateData().

Principal component analysis (PCA) was performed using 30 principal components. Clustering was conducted using the Louvain algorithm at a resolution of 0.3, and two-dimensional visualization was generated using Uniform Manifold Approximation and Projection (UMAP). Differentially expressed genes (DEGs) were identified using FindAllMarkers() with min.pct = 0.01 and logfc.threshold = 0.25.

### Cell Type Annotation and Marker Gene Identification

Cluster-specific marker genes were compared with *Arabidopsis* root markers using the Orthologous Marker Gene Group (OMG) framework [14] (https://github.com/LiLabAtVT/OrthoMarkerGeneGroups). Feature and dot plots were generated using FeaturePlot() and DotPlot() in Seurat.

### Functional Enrichment and Hormone Module Analysis

GO enrichment analysis of cluster-specific marker genes and DEGs between infected and non-infected roots was performed using clusterProfiler v4.14.6 [22], with Benjamini– Hochberg false discovery rate (FDR) correction (adjusted p < 0.05). Hormone-responsive pathway activity was quantified using AddModuleScore() in Seurat. Statistical significance between conditions was assessed using Welch’s t-test with FDR correction. Data visualization was performed using ggplot2.

### Data visualization and reproducibility

All figures were produced in R using Seurat v5.3.0, ggplot2 V4.0.0, and ComplexHeatmap v2.22.0. Scripts and parameters used for preprocessing, integration, and visualization are available upon request.

## Data Availability / Accession Numbers

The raw and processed single-nucleus RNA sequencing (snRNA-seq) data generated in this study have been deposited in the NCBI Sequence Read Archive (SRA) under the BioProject accession number PRJNA1334526.

## Code and Materials Availability

All scripts used for data preprocessing, Seurat analysis (v5.3.0), hormone module scoring, and Gene Ontology enrichment are available at the project’s forthcoming GitHub repository (https://github.com/LiLabAtVT/OomyceteRoot). The repository will include R scripts, parameter settings, and workflow documentation corresponding to the analysis pipeline illustrated in **Supplementary Figure S2**. Any additional materials, such as reference files and curated gene lists for SA, ET, and JA modules, will also be made available through the GitHub repository and upon reasonable request from the corresponding author.

## Supplemental Data

**Supplementary Figure S1**. Nuclei quality assessment and sequencing metrics for single-nucleus RNA-seq datasets.

**Supplementary Figure S2**. Cell type annotation.

**Supplementary Figure S3**. Cell type–specific marker expression and GO enrichment in *Arabidopsis thaliana* root nuclei during *P. capsici* infection.

**Supplementary Figure S4**. Workflow for separating and processing plant and pathogen reads from single-nucleus RNA-seq data.

**Supplementary Dataset S1**: R scripts and metadata for Seurat analysis, hormone module scoring, and GO enrichment are available at the following GitHub repository: https://github.com/LiLabAtVT/OomyceteRoot.git

## Author Contributions

J.M. conceptualized the study, designed the experiment, and funded the research. S.L. and J.M. supervised the project. R.A. analyzed the data and generated the single-nucleus RNA-seq data. T.N.C. performed data analysis. J.S. and R.A. conducted the wet-bench experiments. B.O.R.B. contributed to experimental design and data interpretation. All authors except J.M. contributed to writing the manuscript and approved the final version.

## Acknowledgments

This project was a collaborative effort with Dr. John McDowell from 2022 to 2024. Prior to his passing in 2024, Dr. McDowell conceptualized the study, designed the experiments, and provided financial support to carry out the single-nucleus sequencing work. We thank Novogene for performing the single-nucleus RNA sequencing and for their technical support. We are grateful to members of the Song Li laboratory for helpful discussions and feedback throughout the project. Computational analyses were performed using high-performance computing resources provided by Virginia Tech. We also acknowledge the support of collaborators and colleagues who contributed to experimental design and data interpretation.

---

**Supplementary Figure 1. Nuclei quality assessment and sequencing metrics for single-nucleus RNA-seq datasets. (a)** Pre-library preparation quality control metrics for isolated nuclei from non-infected and *P. capsici*-infected *Arabidopsis* thaliana roots. Parameters include total nuclei density, live nuclei density, dead cell density, and calculated viability. All samples displayed high viability (>83%), supporting their suitability for single-nucleus RNA-seq library preparation. **(b)** Summary of sequencing and library construction metrics generated using Cell Ranger (v8.0.1). Reported metrics include estimated number of nuclei recovered, mean reads per nucleus, median genes detected per nucleus, fraction of reads associated with nuclei, and total sequencing depth. All samples were processed under the same experimental conditions (8/28/24), ensuring comparability across biological replicates and treatment groups.

**Supplementary Figure 2: Cell type annotation**

Cell type annotation using Orthologous Marker Gene Groups (OMGs). The y-axis represents clusters identified in the snRNA-seq dataset, and the x-axis represents cell type clusters from the OMG reference database. Color intensity indicates the number of shared orthologous marker gene groups (darker green = more shared groups, lighter green = fewer shared groups). Asterisks denote statistical significance (* < 0.05).

**Supplementary Figure 3. Cell type–specific marker expression and GO enrichment in *Arabidopsis thaliana* root nuclei during *P. capsici* infection**.

(a) Dot plot showing the top cell type–specific marker genes identified across the thirteen transcriptionally distinct clusters of *Arabidopsis* root nuclei. Each dot represents the percentage of nuclei expressing the gene (dot size) and the average expression level (color gradient). (b) UMAP feature plots displaying representative marker genes for major root tissues: APL (AT1G79430; phloem companion cells), ANAC033 (AT1G79580; procambium-associated transcription factor), CASP1 (AT2G36100; endodermal marker), and ATEXP7 (AT1G12560; root hair-related expansin). Expression intensity is shown by color scale.

**Supplementary Figure 4: Workflow for separating and processing plant and pathogen reads from single-nucleus RNA-seq data**.

The pipeline outlines the steps used to distinguish host (plant) and pathogen reads following multi-species alignment. Sequencing data are first processed using Cell Ranger Count with multi-species reference genomes. The resulting gem_classification.csv file is then used to identify and separate plant and pathogen barcodes. Next, subset-bam is used to generate species-specific BAM files, which are subsequently converted to FASTQ format using bamtofastq. Finally, each FASTQ file is aligned independently to its corresponding reference genome for downstream analysis.

