## Supplemental Data for "Single nucleus transcriptome analysis of *Arabidopsis thaliana* roots infected with *Phytophthora. capsici*"

**a**

| Sample Name | Date | Nuclei Density ( nuclei /ml) | Live Nuclei density (nuclei/ml) | Dead Cells Density (nuclei/ml) | Viability |
|---|---|---|---|---|---|
| Non_Infected_1 | 8/28/24 | 460,000 | 60,775 | 25,975 | 94.63% |
| Non-Infected_2 | 8/28/24 | 335,000 | 43,550 | 56,750 | 83.06% |
| Infected_1 | 8/28/24 | 370,000 | 48,100 | 23,300 | 93.67% |
| Infected_2 | 8/28/24 | 327,500 | 42,575 | 23.35 | 92.91% |

**b**

| Sample name | Non_Infected_1 | Non-Infected_2 | Infected_1 | Infected_2 |
|---|---|---|---|---|
| Plant Species | Arabidopsis thaliana | Arabidopsis thaliana | Arabidopsis thaliana | Arabidopsis thaliana |
| Experimental Date | 8/28/24 | 8/28/24 | 8/28/24 | 8/28/24 |
| Estimated Number of Cells | 8,016 | 8,010 | 8,000 | 8,000 |
| Mean Reads per cell | 110,382 | 124,296 | 61,091 | 62,500 |
| Median Gene Per cells | 642 | 964 | 456 | 158 |
| Fraction Reads in cells | 54.50% | 43.00% | 93.40% | 95.70% |
| Number of Reads | 884,823,811 | 995,613,403 | 488,728,987 | 500,000,000 |
| Pipeline Version | cellranger-8.0.1 | cellranger-8.0.1 | cellranger-8.0.1 | cellranger-8.0.1 |
| Error | Low Fraction Reads in Cells | Low Fraction Reads in Cells | | |

**Supplementary Figure 1. Nuclei quality assessment and sequencing metrics for single-nucleus RNA-seq datasets.** **(a)** Pre-library preparation quality control metrics for isolated nuclei from non-infected and Phytophthora capsici-infected Arabidopsis thaliana roots. Parameters include total nuclei density, live nuclei density, dead cell density, and calculated viability. All samples displayed high viability (>83%), supporting their suitability for single-nucleus RNA-seq library preparation. **(b)** Summary of sequencing and library construction metrics generated using Cell Ranger (v8.0.1). Reported metrics include estimated number of nuclei recovered, mean reads per nucleus, median genes detected per nucleus, fraction of reads associated with nuclei, and total sequencing depth. All samples were processed under the same experimental conditions (8/28/24), ensuring comparability across biological replicates and treatment groups.

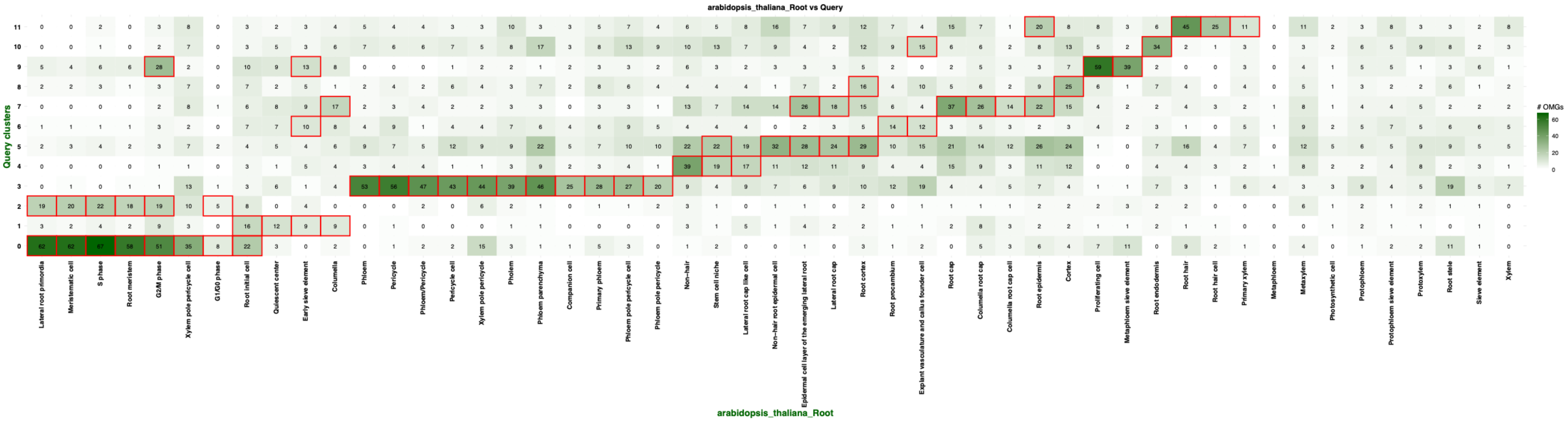


**Supplementary Figure 2: Cell type annotation**

Cell type annotation using Orthologous Marker Gene Groups (OMGs). The y-axis represents clusters identified in the snRNA-seq dataset, and the x-axis represents cell type clusters from the OMG reference database. Color intensity indicates the number of shared orthologous marker gene groups (darker green = more shared groups, lighter green = fewer shared groups). Red boxes denote statistical significance (* < 0.05).

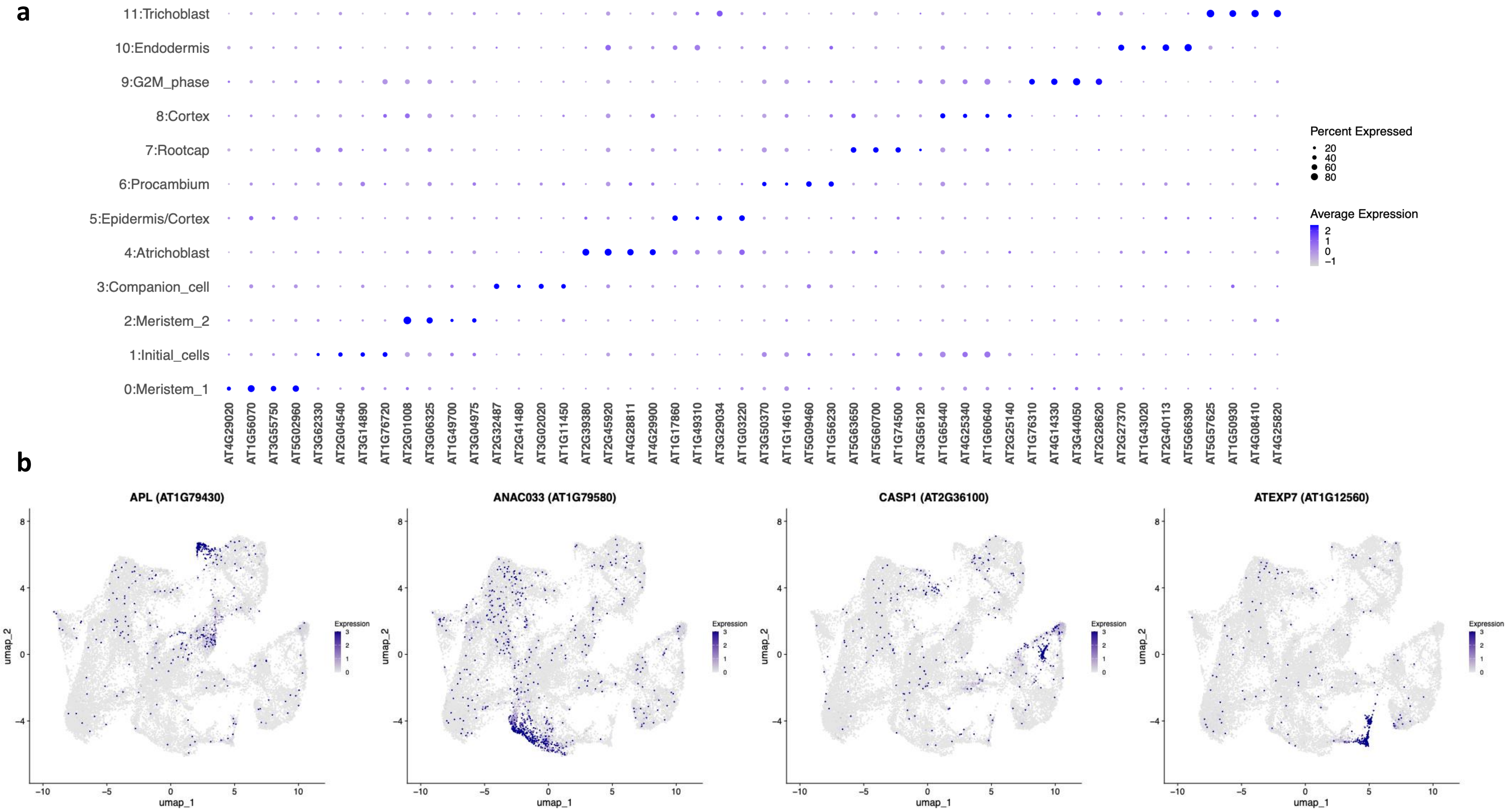
a
11:Trichoblast
10:Endodermis
9:G2M_phase
8:Cortex
7:Rootcap
6:Procambium
5:Epidermis/Cortex
4:Atrichoblast
3:Companion_cell
2:Meristem_2
1:Initial_cells
0:Meristem_1
AT4G29020
AT1G56070
AT3G55750
AT5G02960
AT3G62330
AT2G04540
AT3G14890
AT1G76720
AT2G01008
AT3G06325
AT1G49700
AT3G04975
AT2G32487
AT2G41480
AT3G02020
AT1G11450
AT2G39380
AT2G45920
AT4G28811
AT4G29900
AT1G17860
AT1G49310
AT3G29034
AT1G03220
AT3G50370
AT1G14610
AT5G09460
AT1G56230
AT5G63650
AT5G60700
AT1G74500
AT3G56120
AT1G65440
AT4G25340
AT1G60640
AT2G25140
AT1G76310
AT4G14330
AT3G44050
AT2G28620
AT2G27370
AT1G43020
AT2G40113
AT5G66390
AT5G57625
AT1G50930
AT4G08410
AT4G25820
Percent Expressed
20
40
60
80
Average Expression
2
1
0
−1
b
APL (AT1G79430)
ANAC033 (AT1G79580)
CASP1 (AT2G36100)
ATEXP7 (AT1G12560)
umap_2
umap_1
Expression
−10
−5
0
5
10
−4
0
4
8
3
2
1
0

**Supplementary Figure 3. Cell type–specific marker expression and GO enrichment in Arabidopsis thaliana root nuclei during *Phytophthora capsici* infection.**

**(a)** Dot plot showing the top cell type–specific marker genes identified across the thirteen transcriptionally distinct clusters of Arabidopsis root nuclei. Each dot represents the percentage of nuclei expressing the gene (dot size) and the average expression level (color gradient). **(b)** UMAP feature plots displaying representative marker genes for major root tissues: APL (AT1G79430; phloem companion cells), ANAC033 (AT1G79580; xylem-associated transcription factor), CASP1 (AT2G36100; endodermal marker), and ATEXP7 (AT1G12560; root hair–related expansin). Expression intensity is shown by color scale.

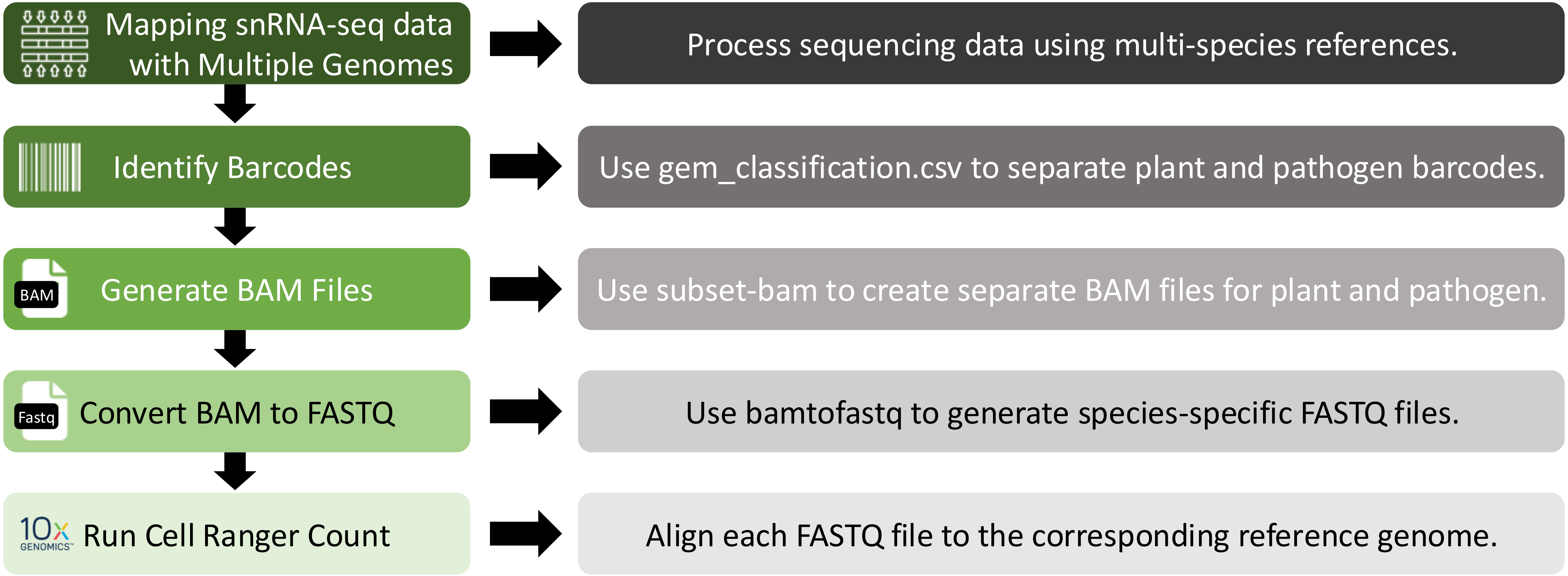


**Supplementary Figure 4:Workflow for separating and processing plant and pathogen reads from single-nucleus RNA-seq data.** The pipeline outlines the steps used to distinguish host (plant) and pathogen reads following multi-species alignment. Sequencing data are first processed using Cell Ranger Count with multi-species reference genomes. The resulting gem_classification.csv file is then used to identify and separate plant and pathogen barcodes. Next, subset-bam is used to generate species-specific BAM files, which are subsequently converted to FASTQ format using bamtofastq. Finally, each FASTQ file is aligned independently to its corresponding reference genome for downstream analysis.
